# Lower-order methylation states underlie the maintenance and re-establishment of Polycomb modifications in *Drosophila* embryogenesis

**DOI:** 10.1101/2025.07.25.666882

**Authors:** Eleanor A. Degen, Natalie Gonzaga-Saavedra, Shelby A. Blythe

## Abstract

Polycomb Group (PcG) proteins regulate the chromatin composition of an embryo by facilitating the mono, di, and tri-methylation of Histone H3 Lysine 27 (H3K27me1/2/3). For the zygote to inherit an H3K27 methylation blueprint from its mother, PcG-modified states established during oogenesis must persist through early embryogenesis until the onset of large-scale zygotic transcription (Zygotic Genome Activation, ZGA). However, questions have persisted regarding the relative contributions of two molecular mechanisms to the propagation of H3K27 methylation through early development: 1) allosteric regulation of the H3K27 methyltransferase Enhancer of Zeste (E(z)) by existing H3K27me2/3, and 2) nucleation of E(z) activity at chromatin by DNA binding factors. Here, we investigate how allostery and nucleation contribute to H3K27 methylation dynamics in early *Drosophila* embryogenesis by developing and experimentally validating a mathematical model. This model incorporates measurements of the nuclear concentration dynamics of E(z) and the Polycomb Response Element binding factor Pleiohomeotic (Pho), as well as the dilution of epigenetic modifications at DNA replication with the incorporation of histones to nascent chromatin. With stochastic simulations and *in vivo* experiments, we assert that allosteric regulation of E(z) maintains a PcG-imprint on maternal chromosomes in the form of lower-order H3K27 methylation states (H3K27me1/2), that de novo establishment of H3K27 methylation at paternal chromosomes relies on nucleation of E(z) activity by Pho, and that broad H3K27me3 domains at both maternal and paternal chromosomes are re-established at ZGA. This work provides a mechanistic explanation for the inheritance of Polycomb states in contexts of intense cellular proliferation.

## INTRODUCTION

Multicellular organisms employ Polycomb group (PcG) proteins during development to establish epigenetic modifications that regulate zygotic gene expression patterns (1–5). The Polycomb Repressive Complex 2 (PRC2) deposits H3K27 mono-, di-, and tri-methylation through its catalytic subunit Enhancer of Zeste (E(z)) (6,7). PRC2 is targeted to genomic loci by DNA binding proteins such as Pleiohomeotic (Pho), and by existing H3K27me2/3 (8–10). In many organisms, changes in histone modification status coincide with cell cycle lengthening and zygotic genome activation (ZGA) (11–17). We have recently shown that, despite high levels of E(z) during early embryonic cell divisions, broad H3K27me3 domains emerge after nuclei import Pho at ZGA (18). This finding raises the question of whether E(z) propagates H3K27 methylation through the cell cycles prior to ZGA, and if so, how.

Prior work has revealed the offspring of different species can inherit epigenetic states from their parents, suggesting that embryos transmit a parental blueprint for H3K27 methylation (19–23). At the onset of *Drosophila* development, H3K27me3 decorates maternally-provided chromatin but is absent from paternal chromosomes (23). This maternal blueprint must persist during the cleavage stage of development despite the unfavorable conditions for PRC2 activity at that time (18). Following fertilization, *Drosophila* embryos undergo a series of thirteen synchronous nuclear divisions, at a frequency of one per 8 to 20 minutes (24). These divisions rapidly dilute epigenetic marks through the incorporation of unmodified histones into chromatin (25). Prior studies have reported conflicting findings regarding whether maternally-inherited H3K27me3 can survive the cleavage divisions (11,15,18,23,25–27). Our recent work shows that H3K27me3 is undetectable on the genome by at least nuclear cycle (NC) 8, and is re-established over the one-hour period of NC14 that coincides with large-scale ZGA (18). In later stage embryos and larval tissues, maintenance of PcG states depends on the function of Polycomb Response Elements (PREs), which are bound by nucleating factors such as Pho (28,29). These nucleating factors are absent from nuclei until shortly before ZGA, suggesting PcG states may not be maintained in early embryos through nucleation of E(z) activity at PREs (18). However, nucleation-independent, allosterically regulated E(z) activity could allow for maintenance of the maternal PcG imprint through lower-order H3K27 methylation states (30,31).

The relative contributions of allostery and nucleation to the maintenance of PcG modifications has been subject to mathematical modeling (25,27). A recent model focusing on early *Drosophila* development predicts the observed loss of H3K27me3, but does not simulate the observed rapid re-establishment of H3K27me3 during NC14 (11,15,18,25,27). Since the formulation of this model, recent work has reported additional measurements that could help improve and evaluate the model’s performance (18,25). Here, we update the stochastic model to account for these measurements. The updated model indicates that early embryos rely on allosteric enhancement of E(z) to propagate a maternal imprint of H3K27me2 in *cis*, while the acquisition of all forms of H3K27 methylation at paternal chromosomes relies on nucleating factors operating in *trans*. We validate this prediction experimentally. Our work indicates that PcG cofactor activity and cell cycle lengthening allow parental asymmetries in modification states to resolve at ZGA.

## RESULTS

### E(z) catalyzes a sequence of reactions to rapidly establish H3K27me3 at PREs

We first sought to determine an appropriate output of PcG activity to model. In recent work, we measured the genome-wide occupancy of E(z), Pho, and H3K27me3 via ChIP-seq, finding that H3K27me3 spreads bidirectionally from sites of nucleated E(z) activity over NC14 (Fig. 1A) (18). After approximately an hour post-mitosis 13, domains of H3K27me3 have formed over PREs associated with E(z) and Pho (Fig. 1A) (18). PREs produce characteristic methylation dynamics that are most easily evident at sites spaced sufficiently far apart along the genome. For well-spaced PREs—such as those associated with *knirps* (*kni*) and *knirps-like* (*knrl*)—we see H3K27me3 first accumulates at mid-NC14, and then spreads from these sites over the remainder of the nuclear cycle (Fig. 1A). The dynamics of the nucleation and spreading of H3K27me3 are less evident at more closely spaced PREs (18). Therefore, we analyzed the activity of E(z) peaks located in isolation to characterize the H3K27 methylation levels that a single PRE generates over time (Fig. 1B-D). We find that, at isolated PREs, H3K27me3 increases in a roughly linear fashion at the location of a central E(z) peak (Fig. 1B, D). These dynamics reflect the upgrading of lower-order methylation states, as H3K27me1 is progressively depleted at isolated PREs with the emergence of H3K27me3 domains (Fig. 1C). We conclude that the methylation states of isolated PREs are informative for modeling the molecular mechanisms that define the dynamics of PcG modifications during early development. Further, the depletion of H3K27me1 throughout NC14 highlights the importance of modeling the H3K27 methylation reaction network for understanding the re-establishment of H3K27me3.

**Figure 1:**
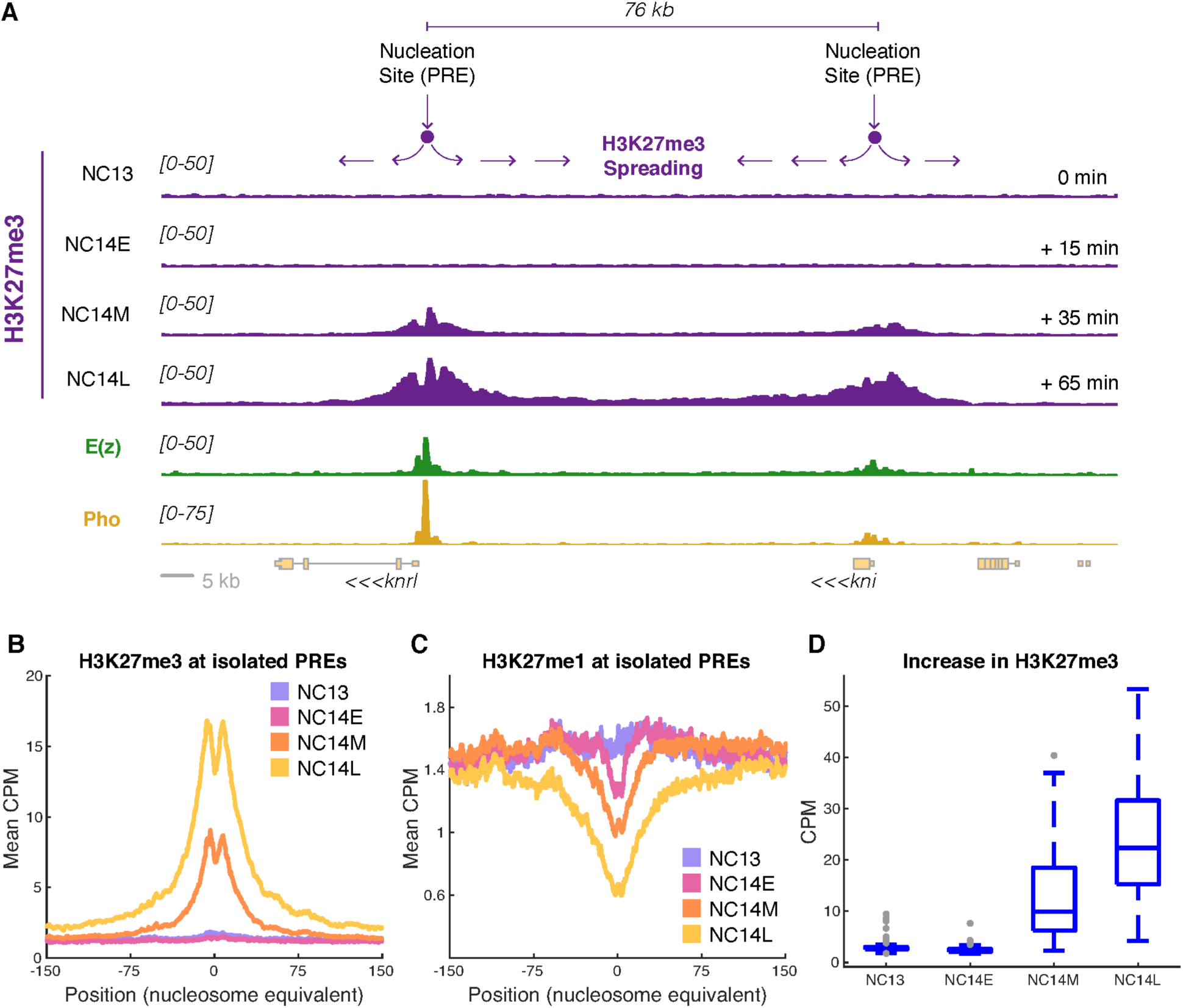
H3K27 methylation accumulates at PREs over NC14. **A)** Shown are the counts-per-million (CPM) normalized ChIP-seq measurements of H3K27me3, E(z), and Pho along a genome track containing the *knrl* and *kni* loci (data from Gonzaga-Saavedra et al., 2025) (18). H3K27me3 was measured in embryos staged at NC13, NC14-early (NC14E), NC14-mid (NC14M), and NC14-late (NC14L) (18). Y-axis CPM ranges are indicated in brackets at left. E(z) and Pho occupy nucleation sites (PREs, annotated at top), around which H3K27me3 spreads throughout NC14. At this region, PREs are spaced sufficiently far apart such that there is not substantial merging of H3K27me3 domains by NC14L. Scale bar = 5 kb. **B)** Analysis of mean H3K27me3 signal flanking isolated E(z) peaks as measured via ChIP-seq at NC13, NC14E, NC14M, and NC14L time points. X-axis values are reported in nucleosome equivalents, with one nucleosome equal to 180 bp. H3K27me3 increases throughout NC14. **C)** Analysis of mean H3K27me1 signal flanking isolated E(z) peaks as measured via ChIP-seq at NC13, NC14E, NC14M, and NC14L time points. As in B, X-axis values are reported in nucleosome equivalents. H3K27me1 is progressively depleted with the formation of H3K27me3 domains. **D)** Analysis of H3K27me3 ChIP-seq signal flanking 128 isolated E(z) peaks. Plotted are distributions of the 99th percentiles of the H3K27me3 CPM measurements associated with the isolated PREs at each measured time point (NC13, NC14E, NC14M, NC14L). H3K27me3 increases in a linear fashion between NC14E and NC14L.

### A revised stochastic approach for modeling H3K27 methylation dynamics

We computationally modeled H3K27 methylation across cell divisions given allosteric regulation of E(z) by existing H3K27me2/3, as well as nucleation of E(z) activity by the DNA binding factor Pho (Fig. 2A, Methods) (10,30). Unlike prior work, our model allows E(z) to methylate H3K27 only if either allostery or nucleation occurs (Fig. 2A, Methods) (25). For allosteric stimulation of E(z), at least one of the modeled histones neighboring a nucleosome of interest must be H3K27me2/3, and E(z) must be associated with the modeled PcG domain. For nucleation, an allostery-driven reaction must not have occurred and both E(z) and Pho must be simultaneously associated with the PcG domain. To implement these mechanisms, we incorporated estimated Pho and E(z) concentrations based on live imaging measurements into the stochastic modeling framework (Fig. 1B, Fig. S1, Methods) (18). We suspected that alterations to the model would allow it to predict substantial levels of H3K27me3 during NC14 given a similar reaction probability to prior work (25). Therefore, we varied the value of the model’s baseline methylation probability (*V*) from approximately the value used in Lundkvist et al. (25), *V* = 5x10^-4^, to *V* = 2x10^-3^ in steps of *V* = 2.5x10^-5^, and evaluated the model’s performance at these values. We find that the sum of the differences between the simulated and measured normalized H3K27me3 values at each time point reaches a minimum at *V* = 1.075x10^-3^ (Fig. 1C), 1.72x the baseline methylation probability of Lundkvist et al. (25). We therefore used *V* = 1.075 x10^-3^ as the baseline methylation probability in all the simulations presented in this work.

**Figure 2:**
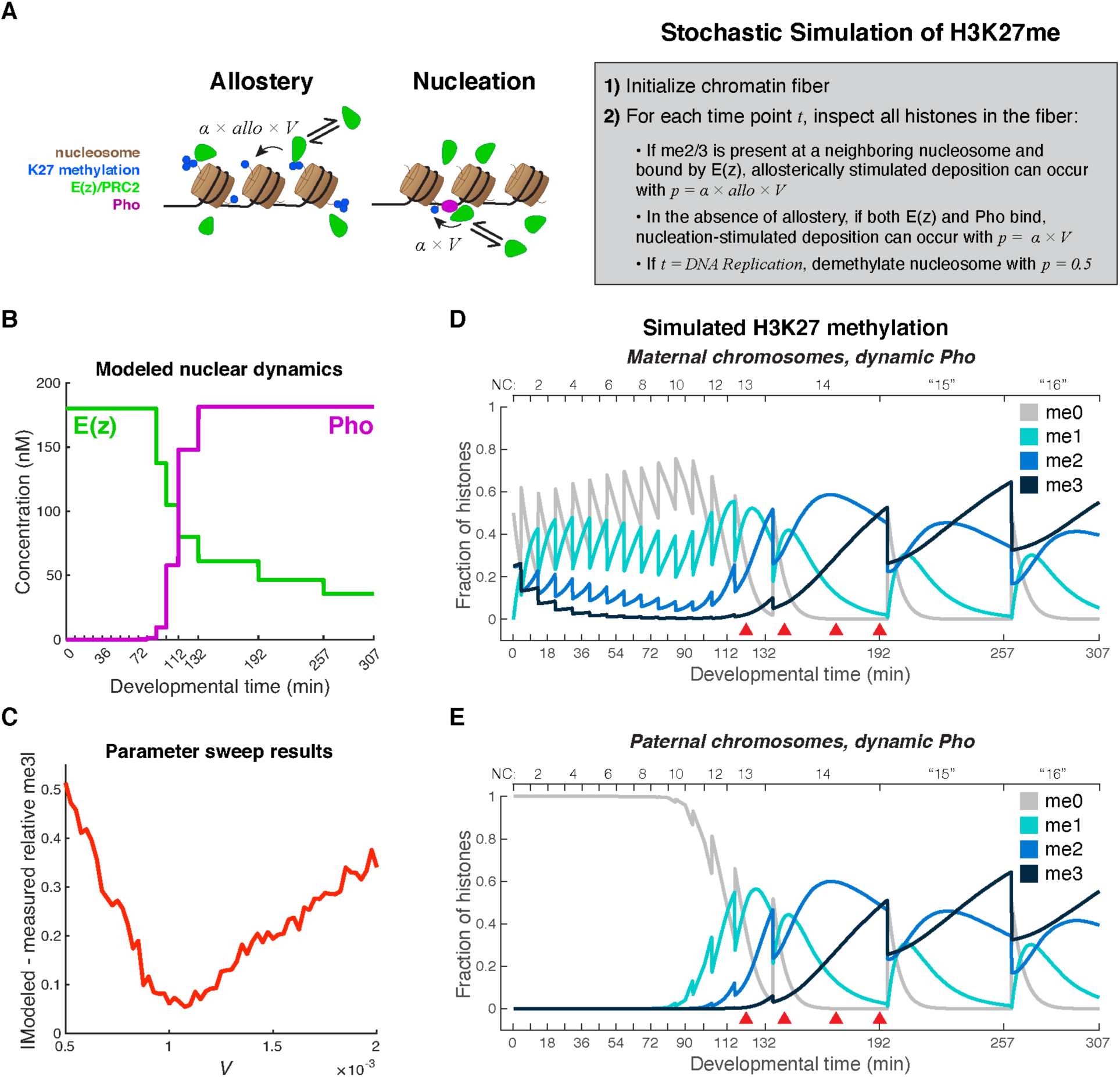
Stochastic simulation of H3K27 methylation dynamics. **A)** Schematic of the simulation strategy. In the model, E(z) (PRC2, green) can methylate a nucleosome within an array if allosterically stimulated by existing H3K27me2/3, or if Pho has concurrently bound the locus (diagram at left). Methylation reactions are modeled as a series of stochastic decisions (outlined at right). At *t* = 0, a chromatin fiber is initialized with an initial H3K27 methylation composition. At each subsequent timepoint, the algorithm inspects the nucleosome array to determine whether methylation groups are added (when successful allostery- or nucleation-driven reactions occur), or removed (at the timepoints of DNA replication). The simulation strategy is outlined in detail in the Methods and is a modified form of a recently published algorithm (25). In the schematic, α is a constant that scales the baseline methylation probability, *V*, according to the methylation substrate (α = 14 for H3K27me0, 4 for H3K27me1, and 0 for H3K27me3). The parameter *allo* and acts to scale the methylation probability when allosteric stimulation of E(z) occurs. **B)** Modeled nuclear concentration dynamics of E(z) and Pho based on prior live imaging measurements of EGFP-E(z) and Pho-sfGFP (Methods) (18). **C)** Shown are the results of a parameter sweep for the baseline methylation probability, *V*, from 5x10^-4^ to 2x10^-3^ in steps of 2.5x10^-5^, holding all other parameters constant (*allo* = 3.5, *K_e_* = *K_p_* = 10 nM, Methods). The absolute value of the sum of the differences between the predicted and measured relative H3K27me3 levels at NC14-early, NC14-mid, and NC14-late are shown for the tested values of *V*. *V* = 1.075 x10^-3^ best predicts the NC14 H3K27me3 relative ChIP-seq values. **D-E)** Modeled dynamics of H3K27 methylation over the first 307 minutes of development (16 cell divisions, Methods). In each plot, shown are the mean fractions of H3K27 positions in a 300-nucleosome chromatin fiber with me0, me2, and me3 marks calculated from n = 100 simulations. For maternal chromosomes, simulations were performed with the initial condition where 50% of H3K27 positions were me0, 0% were me1, 25% were me2, and 25% were me3. For paternal chromosomes, simulations were performed with the initial condition where 100% of H3K27 positions were me0. Red arrowheads mark the time points when H3K27me3 was measured via ChIP-seq (NC13 = mitosis 12 + 10’, NC14E = mitosis 13 +15’, NC14M = mitosis 13 + 35’, NC14L = mitosis 13 + 60’).

We find that, following parameter fitting to the *in vivo* data, our model predicts measured H3K27me3 dynamics at isolated PREs given either maternal or paternal initial H3K27me conditions (Fig. 2D-E). Consistent with our measurements, simulated PcG domains begin NC14 with nearly no H3K27me3 and acquire the mark in a linear fashion over the nuclear cycle (Fig. 2D-E). In contrast to prior work, our simulations predict substantial H3K27me3 levels by the end of NC14 (Fig. 2D-E) (25). The model’s simulation of H3K27me3 domain formation over approximately an hour of development stems from the modeled ability of any directly neighboring H3K27me2/3 groups to allosterically enhance the E(z) activity, as well as the model’s adjusted baseline H3K27me2-me3 reaction probability (25). Further, our model predicts that multiple E(z) molecules simultaneously act at a locus; between the maternal and paternal simulated conditions, on average, 17 E(z) molecules complete a methylation reaction per minute. This suggests that, by operating in collectives of molecules that can be allosterically stimulated by existing H3K27me2/3, E(z) can overcome a relatively slow catalytic rate to establish broad PcG domains over a short time period.

While the model predicts similar H3K27me3 dynamics at maternal and paternal chromosomes over NC14, it simulates notable differences in their H3K27 methylation composition earlier in development. If the model incorporates initial conditions that describe maternal chromosomes— which begin development with H3K27me2/3—it simulates dilution of H3K27me3 with each round of genome replication until approximately NC6, consistent with prior work (Fig. 1D) (25). In addition, with maternal initial conditions, the model predicts that E(z) successfully maintains substantial levels of H3K27me1/2 over the early nuclear cycles (Fig. 1D). In contrast, the model does not predict any form of H3K27 methylation at paternal chromosomes—which begin development without H3K27 methylation (23)—until H3K27me1 begins to accumulate at NC9 (Fig. 1E). H3K27me2 does not begin to be simulated at paternal chromosomes until NC12 (Fig. 1E). The differences between the maternal and paternal simulations suggest that they may rely differentially on the mechanisms that allow for H3K27 methylation: allosteric stimulation of E(z) by existing H3K27me2/3 vs. nucleation of E(z) activity by a DNA binding factor such as Pho.

### Nucleating factors contribute to the parent-of-origin asymmetry of Polycomb modifications

We used the model to test whether H3K27 methylation at maternal and/or paternal chromosomes is limited by the absence of nucleating factors in nuclei prior to NC11 (18). If the nuclear dynamics of DNA binding factors limit H3K27me3 prior to NC14, then altering the model to incorporate a constant high nuclear Pho concentration would lead to higher simulated levels of H3K27me3 during that time. Simulations show a constant nuclear Pho concentration does not dramatically change the modeled dynamics of H3K27me3 at maternal or paternal chromosomes; significant H3K27me3 deposition is delayed until NC14 under any condition (Fig. 3A-B, Fig. 2D-E). However, constant high nuclear Pho concentrations do produce a dramatic shift in the simulated H3K27me1/2 dynamics at paternal chromosomes (Fig. 3B, Fig. 2E). With abundant nuclear Pho throughout the early nuclear cycles, the model now predicts substantial H3K27me1/2 at both maternal and paternal chromosomes during this time (Fig. 3A-B). The contrast between the constant Pho and dynamic Pho simulations suggests that the absence of nucleating factors prevents the acquisition of H3K27 methylation at paternal chromosomes prior to NC9. We expect the reliance of the paternal chromosomes on nucleation delays them from “catching up” to the PcG state of the maternal chromosomes. If simulated nuclear Pho levels are constant, the difference between the predicted mean paternal and maternal methylation states reaches zero around NC6. In contrast, when simulated nuclear Pho is dynamic, predicted paternal and maternal methylation levels do not match until mid NC14 (Fig. 3C). Our data supports the conclusion that allosteric regulation allows the maternal PcG state to propagate through the early cell cycles independently of nucleation, while the acquisition of H3K27 methylation on the paternal chromosomes is limited by the availability of nucleating factors.

**Figure 3:**
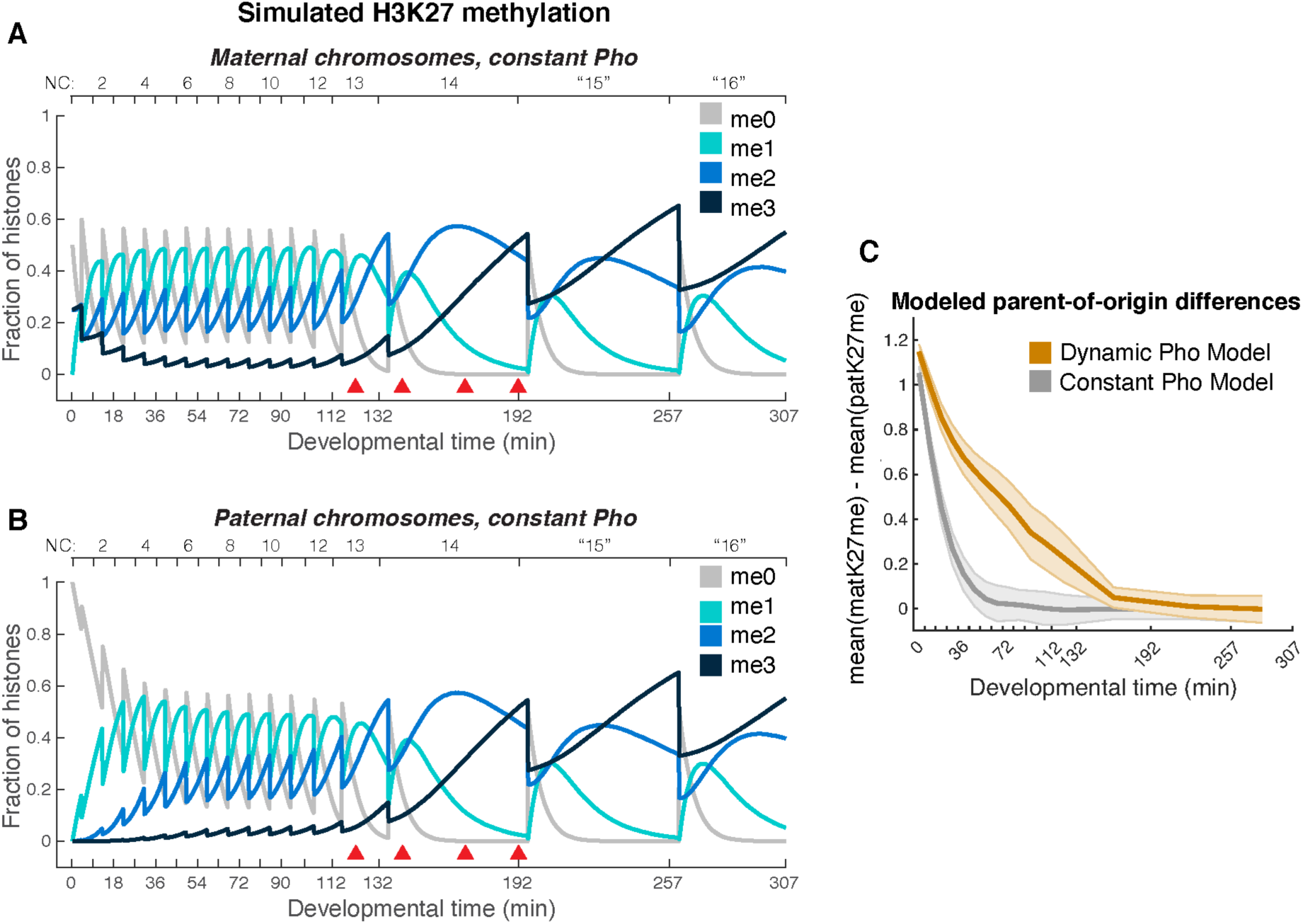
Modeled parent-of-origin differences. **A-B)** Modeled dynamics of H3K27 methylation over the first 307 minutes of development (16 cell divisions, Methods) when nuclear Pho is held at a constant maximal value (181 nM). In each plot, shown are the mean fractions of H3K27 positions in a 300-nucleosome chromatin fiber with me0, me2, and me3 marks calculated from n = 100 simulations. For maternal chromosomes, simulations were performed with the initial condition where 50% of H3K27 positions were me0, 0% were me1, 25% were me2, and 25% were me3. For paternal chromosomes, simulations were performed with the initial condition where 100% of H3K27 positions were me0. **C)** Shown are the differences between the mean methylation states of maternal and paternal chromosomes as calculated on a per-nuclear cycle basis for the Dynamic Pho (Fig. 2 D-E) and Constant Pho simulations (Fig. 3 A-B). If nuclear Pho concentrations are limited in the early nuclear cycles, paternal chromosomes take longer to catch up to the PcG state of the maternal chromosomes than when nuclear Pho concentrations are high from the beginning of development.

### H3K27me2 is propagated on maternal chromosomes from the beginning of development

Our model predicts that maternal chromosomes maintain lower-order H3K27 methylation throughout the earliest nuclear divisions, while paternally-derived chromosomes do not (Fig. 2D-E). We sought to experimentally test this prediction by immunostaining H3K27me2 in early wild type embryos and in haploid embryos lacking paternal chromosomes (Fig. 4). Our measurements in wild type embryos reveal extensive maintenance of an H3K27me2 modified state (Fig. 4A). Repeating the H3K27me2 staining in *E(z)* mutant embryos yields no measurable signal, confirming that the signal detected by this approach is specific to PcG activity (Fig. 4B). Similar to -me3, H3K27me2 staining of nuclei shortly after fertilization decorates only a portion of chromatin in each nucleus (Fig. 4A, NC2) (18,23). Unlike -me3, however, H3K27me2 staining remains detectable within nuclei throughout the early nuclear divisions (Fig. 4A, NC6, NC13, & Fig. 4C) before increasing in intensity at NC14 (Fig. 4A, NC14) (18). We noted that, in all of the pre-ZGA specimens, the H3K27me2 signal marked only a subregion of the nucleus, similar to H3K27me2 and -me3 staining near the time of fertilization (18,23). To better visualize this, we identified pre-NC14 embryos at mitotic prophase and generated high-resolution images. In all cases, when individual chromosomes become discernible at prophase, the high-resolution H3K27me2 signal shows a punctate distribution on only a subset (approximately 50%) of DAPI-positive material (Fig. 4D-E). We hypothesized that this partial staining pattern reflects maintenance of the H3K27me2 modification on maternal chromatin.

**Figure 4:**
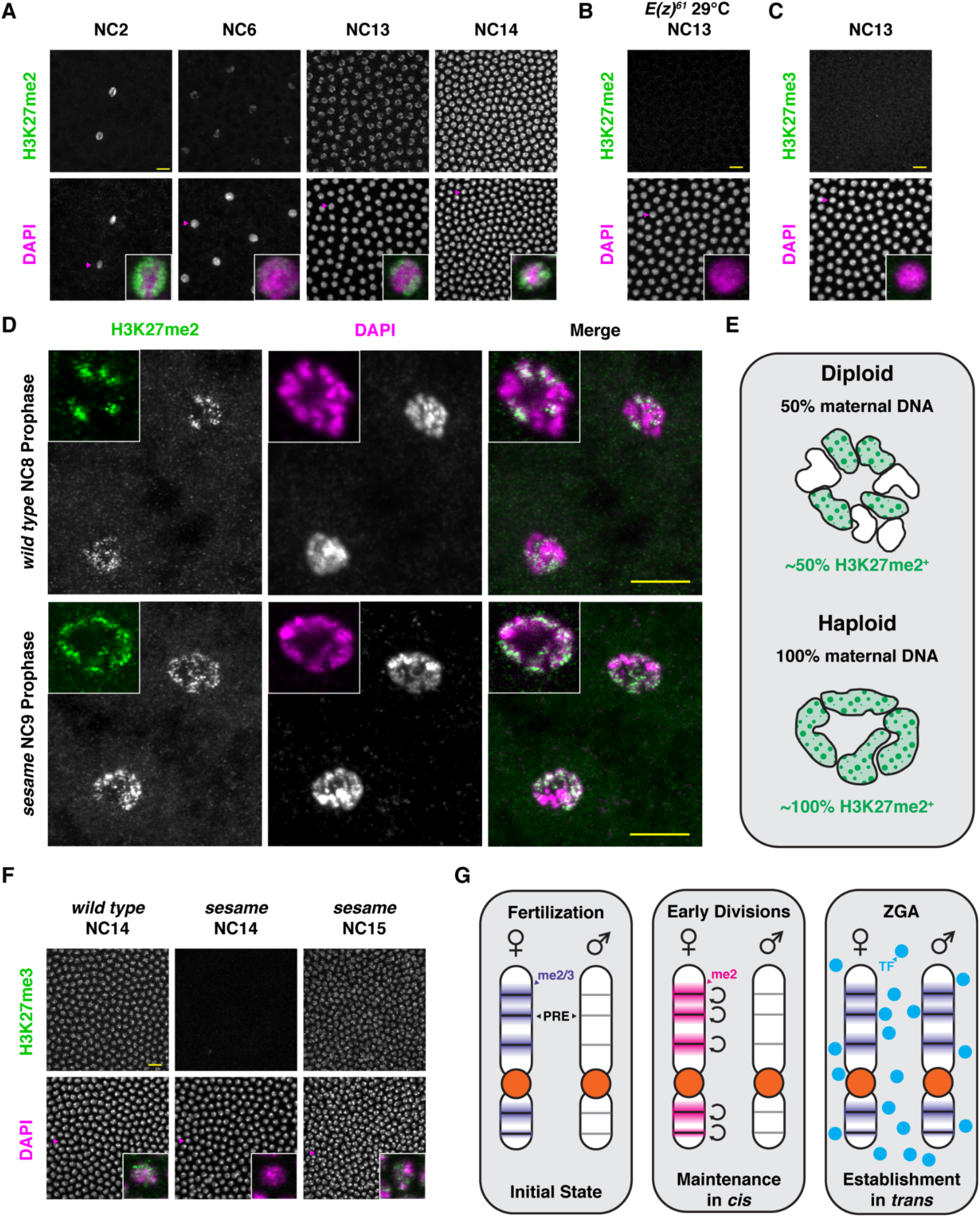
H3K27me2 is propagated on maternal chromosomes in *cis* throughout the cleavage divisions. **A)** H3K27me2 is detected on chromatin throughout early development. Representative images are shown for early embryos at the indicated stages immunostained for H3K27me2 (top) and co-stained for DAPI (bottom). Merged images for representative nuclei (magenta arrowheads) for each stage are shown in the insets (green = H3K27me2, magenta = DAPI). Image intensities are scaled to allow direct quantitative comparison of staining intensity between images. Scale bar (yellow) is 10 µm. **B)** Early embryonic H3K27me2 deposition depends on *E(z)* function. Shown is a representative image of an NC13 embryo from an *E(z)^61^* mother collected at 29°C stained for H3K27me2 and DAPI and displayed as in panel A. Scale bar (yellow) is 10 µm. **C)** H3K27me3 is not strongly detected on NC13 chromatin. Shown is a representative image of an NC13 embryo from a wild type mother stained for H3K27me3 and DAPI and displayed as in panel A. Scale bar (yellow) is 10 µm. **D)** H3K27me2 marks maternally derived chromatin. Wild type (top) or *sesame* mutant (bottom) embryos were stained for H3K27me2 and DAPI. The images show maximum projections of two nuclei from representative mid-cleavage stage embryos at prophase of the cell cycle. Insets show a single z-plane bisecting a single nucleus to highlight condensing chromosome figures. While a subset of DAPI-positive material in wild type embryos is stained for H3K27me2, all DAPI-positive material in *sesame* mutant embryos show punctate H3K27me2 staining. Scale bar (yellow) is 10 µm. **E)** Cartoon representation of insets in panel D. DAPI staining for wild type and *sesame* mutant was traced and color coded to indicate scored H3K27me2 positive material. Haploid embryos from *sesame* mutants only contain maternally-derived chromatin. **F)** Delayed cell cycle lengthening delays deposition of H3K27me3. Wild type (left) or *sesame* mutant (middle and right) embryos were stained for H3K27me3 and DAPI and displayed as in panel C. H3K27me3 accumulates on chromatin at NC14 in wild type embryos (left). *sesame* mutants undergo an additional division before undergoing cell cycle lengthening. H3K27me3 is absent on *sesame* chromatin at NC14 (middle), but accumulates to comparable levels one cell cycle later at NC15 (right). Scale bar (yellow) is 10 µm. **G)** Cartoon summary of H3K27 methylation dynamics in early *Drosophila* development. The maternal genome is marked by H3K27me2/3 and this mark is absent from the paternal genome (left). During the cleavage divisions, the maternal imprint is maintained *in cis* at the level of H3K27me2 (middle). Once *trans-*acting factors gain nuclear localization during ZGA, they operate to re-establish methylation patterns on paternal chromatin (right).

We tested the hypothesis that H3K27me2 decorates maternal rather than paternal chromosomes during the cleavage divisions by staining haploid embryos produced by *sesame* mutant mothers (Flybase: *hira*). *sesame* mutants develop as gynogenetic haploids due to failure of male pronuclear decondensation following fertilization (32,33) and therefore alter the early developmental environment in two key ways relevant to our model. First, *sesame* zygotic nuclei only contain maternally derived chromatin. In contrast to patterns in wild type embryos, the punctate distribution of H3K27me2 staining covers nearly all of the chromatin mass in *sesame* mutant nuclei (Fig. 4D-E). This supports the hypothesis that H3K27me2 is limited to maternal chromatin in wild type embryos. Second, haploid embryos undergo an additional nuclear division to reach the nucleocytoplasmic ratio necessary to trigger cell cycle lengthening and ZGA (34,35). While wild type embryos begin to accrue H3K27me3 following cell cycle lengthening at NC14, *sesame* mutants delay re-establishment of H3K27me3 until NC15 (Fig. 4F). This supports a prediction from our stochastic simulation: the rapid pace of the cleavage divisions limits H3K27me3 deposition in early embryos. Overall, we conclude that maintenance and re-establishment of PcG modification states during early development is constrained by cell cycle activity, and that a maternal PcG imprint is maintained *in cis* on chromatin primarily in the form of H3K27me2 (Fig. 4G).

## DISCUSSION

Here, we show that, during early *Drosophila* development, the parent-of-origin asymmetries in H3K27 methylation state are maintained *in cis* at the level of H3K27me2 by allosteric stimulation of E(z)/PRC2 activity. This activity is not predicted from rate measurements and *in vivo* observations from cell culture systems, suggesting that PRC2 in early *Drosophila* embryos operates with relatively enhanced catalytic throughput. However, despite a likely enhanced E(z) catalytic rate, the pace of the early embryonic cleavage divisions limits the emergence of large tracts of H3K27me3 until cell cycle lengthening at ZGA. In support of this conclusion, our computational model predicts an absence of substantial H3K27me3 on cleavage-stage chromatin, ChIP and immunofluorescence measurements do not show robust levels of H3K27me3 on chromatin prior to NC14, and delaying cell cycle lengthening by one division correspondingly delays high-level acquisition of H3K27me3. As embryos undergo ZGA, PRE-binding factors such as Pho localize to nuclei and nucleate genome-wide patterns of H3K27 methylation, including the *de novo* modification of the paternal genome. We predict that parent-of-origin asymmetries are effectively equalized by late NC14 as embryos engage in pattern formation and cell fate specification.

How are high methylation levels achieved in early *Drosophila* development despite low measured catalytic rates for E(z)? While *in vivo* measurements of mammalian orthologs of E(z) reflect catalytic rates on the order of tens of hours, *Drosophila* embryos establish robust tracts of H3K27me3 modified chromatin on the order of an hour (15,18,31,36,37). Our computational model simulates, on average, 4.2 successful methylation reactions per 15-second time step, suggesting that E(z) operates in clusters of approximately 17 molecules, as E(z) has a refractory period of at most one minute after catalyzing a reaction (38–40). While EGFP-tagged E(z) does not apparently form large high-concentration assemblies at early developmental stages as expected for Polycomb Repressive Complex 1 (PRC1) (18,41–43), mass spectrometry studies indicate that PRC2 components are maintained at high bulk concentrations in early embryos (44). Taken together, these results suggest that PRC2 collectives operating at high concentrations may be key for the maintenance of the maternal PcG imprint and rapid re-establishment of PcG domains at ZGA. Additionally, the chromatin state of early embryos may favor rapid E(z) catalysis. While benchmarking for PRC2 catalysis has been based on mammalian cells otherwise at equilibrium for a specific epigenetic state, such states are nascent in early *Drosophila* embryos (36,39). For instance, early embryos lack Histone H3 Lysine 36 methylation, a mark that antagonizes H3K27me3 deposition (15,45). The early *Drosophila* embryo may therefore provide a more direct experimental system than cultured cell models to evaluate the mechanism for establishment of epigenetic states associated with the Polycomb system.

Our computational model predicts that, despite the absence of high-level H3K27me3 on cleavage-stage chromatin, the embryo propagates the maternal PcG imprint through lower-order H3K27 methylation states. We validate this prediction experimentally, observing persistent H3K27me2 on the ∼50% of chromatin that corresponds to the maternal genome where the modification is epigenetically maintained *in cis*. Notably, maternal PcG imprints are also propagated through H3K27me2 during mouse embryogenesis (46). In the typical cell, H3K27me1/2 modifications are broadly distributed across most of the genome, largely outside of PcG domains, with some enrichment for loci with low transcriptional activity (47,48). Such broad domains of H3K27me2 are thought to stem from a transient and perhaps nonspecific “hit-and-run” mode of deposition distinct from nucleated deposition at PcG domains (47–49). We favor an interpretation where the punctate H3K27me2 signal we observe on cleavage stage chromatin reflects incomplete maturation of the K27 methyl-series at PcG domains, as opposed to genome-wide modifications deposited by a non-specific “hit-and-run” mechanism. Further, the lack of H3K27me2 staining on paternal chromatin during cleavage stages reflects the absence of *trans*- acting factors in early cleavage divisions. It also suggests that ‘hit-and-run’ mechanisms do not have a substantial role during early cleavages. Future epigenomic studies will address the genomic distribution of H3K27 modifications as a function of developmental progression.

Compared with -me3, the H3K27me2 modification is less well understood both in terms of its role in gene regulation, as well as how it could direct allosteric regulation of PRC2. As the observed H3K27me2 signal likely reflects incomplete maturation of the K27 methyl-series at PcG domains, we propose two possible mechanisms for allosteric maintenance of the maternal PcG state *in cis*. In one case, additional factors could allosterically stimulate E(z) by “reading” the H3K27me2 modification. Alternatively, trace amounts of H3K27me3 remaining following DNA replication could allosterically stimulate maintenance of the maternal PcG state. Additional work is needed to distinguish these possibilities. We note that the key allosteric regulatory subunit for PRC2, the *Drosophila* Extra sex combs (Esc) or mammalian Embryonic Ectoderm Development, shows affinity for the -me2 state, although it preferentially binds H3K27me3 (50–53). Further, additional H3K27me2 binding factors have been identified through affinity purification strategies (52), and orthologs for several of these are maternally expressed in *Drosophila.* Whether the mechanism for allosteric regulation by H3K27me2 depends on Esc or dedicated -me2 reading modules represents an area for future investigation.

## MATERIALS AND METHODS

### 1. Stochastic simulations

#### 1.1 Algorithm

We modified the computational Monte Carlo model presented in Lundkvist et al. to simulate the dynamics of H3K27 methylation along a chromatin fiber (25). In this algorithm, a chromatin fiber is represented by an array of nucleosomes, each with two H3K27 methylation positions. As in prior work, at 15 second intervals, the algorithm inspects the methylation state of each position and evaluates whether the position gains or loses a methylation group in that time interval (25). Our model employs an allosteric constant of *allo* = 3.5, equivalent to a prior allosteric parameter value, to scale the probability E(z) completes a methylation reaction if any of the H3K27 positions neighboring a nucleosome of interest contain me2/3 (Section 3.2: Allostery) (25). Our model provides E(z) the opportunity to methylate if allosteric stimulation occurs, or if the nucleating factor Pho binds to the simulated PcG domain along with E(z) (Section 3.2: Allostery, Section 3.3: Nucleation). Unlike prior work, we do not model “hit-and-run” E(z) activity throughout the genome (25). Instead, we model the methylation dynamics of a prospective PcG domain containing a PRE that can be occupied by both E(z) and Pho. Below we present the structure of the stimulation method.

Initialize a 300-nucleosome chromatin fiber (Section 1.2: Initial conditions)

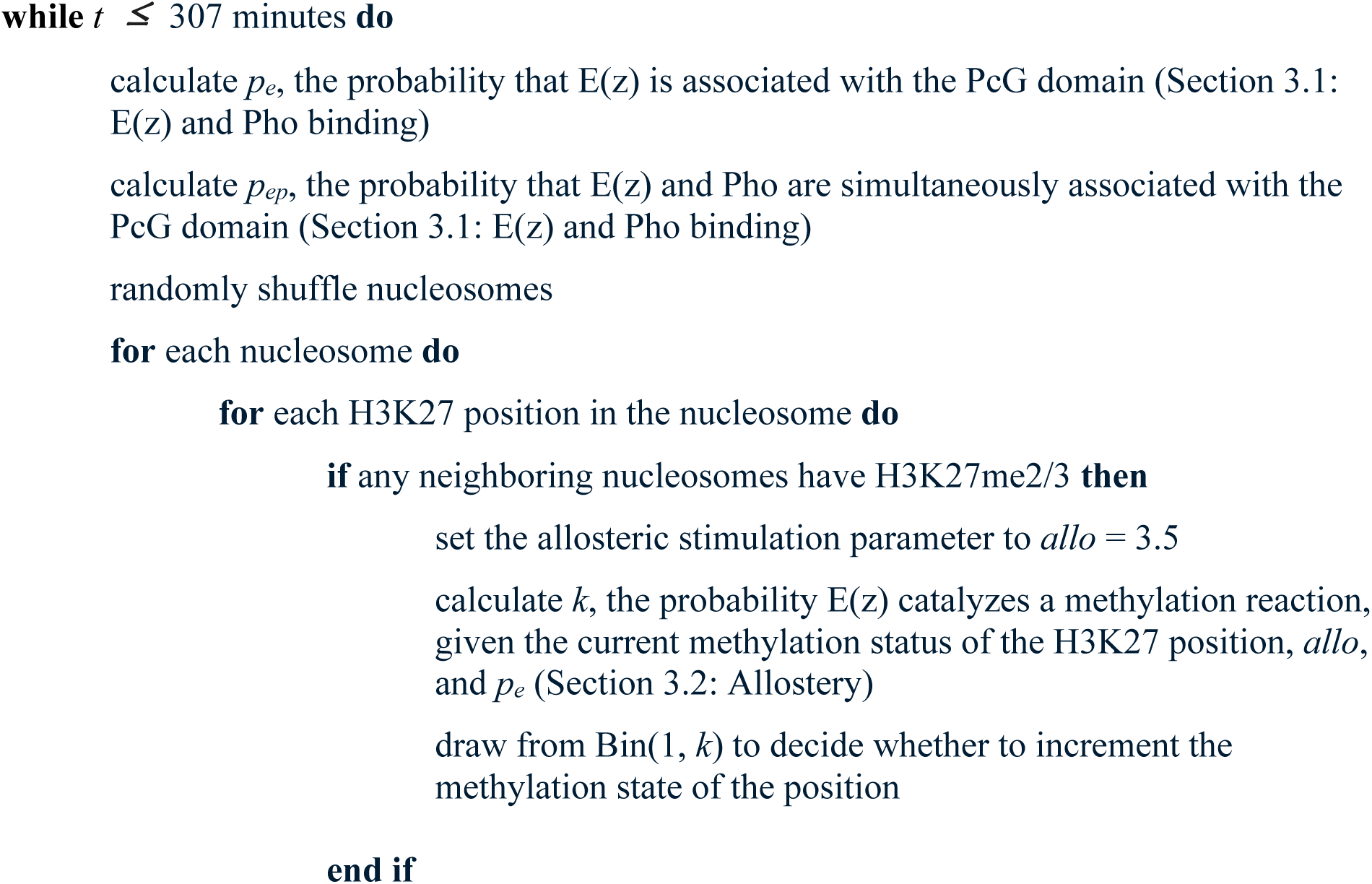
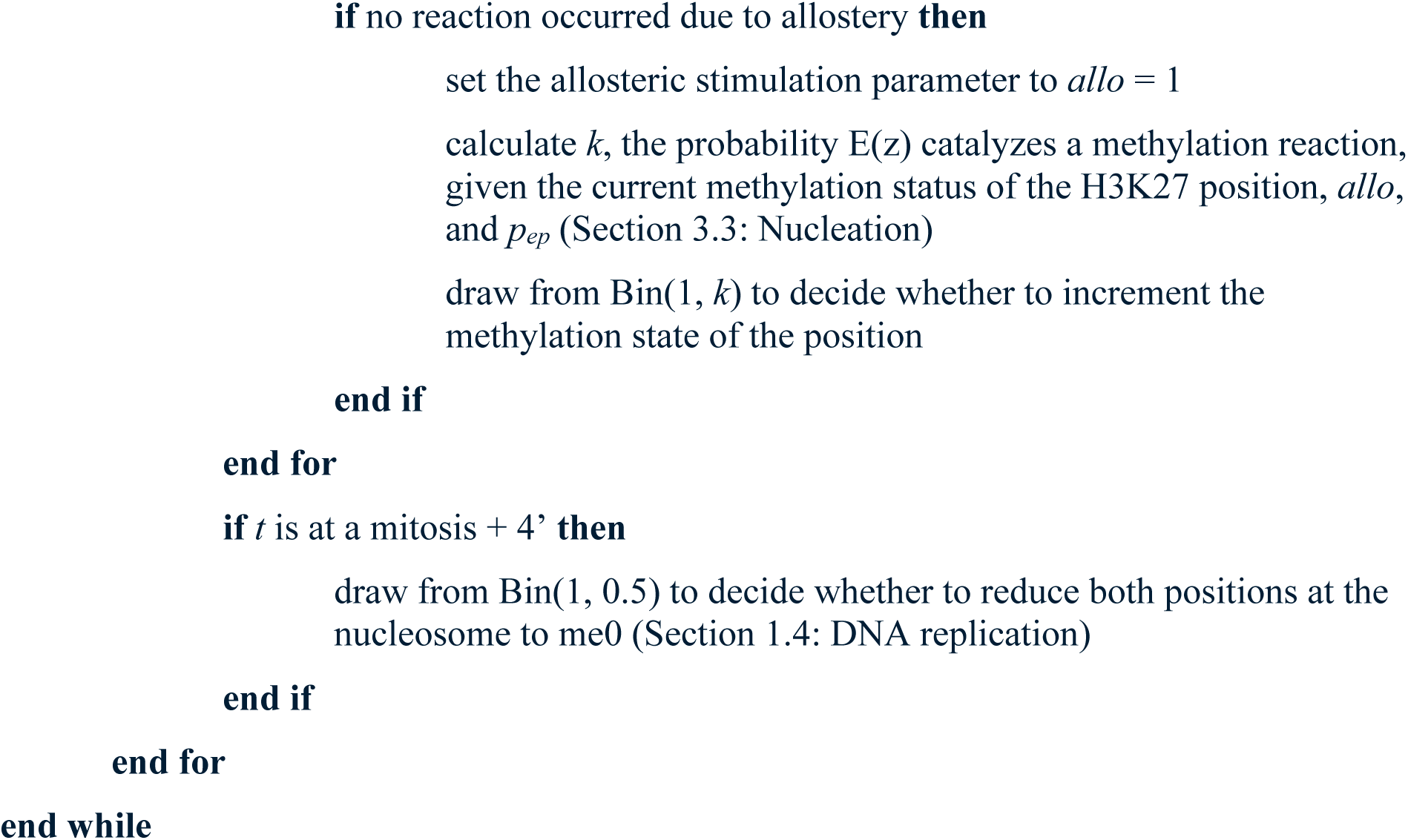

#### 1.2 Initial conditions

To simulate maternally-provided chromatin, in each simulation a chromatin fiber was initialized with 50% of its H3K27 positions with me0, 0% of its positions with me1, 25% of its positions with me2, and 25% of its positions with me3. Our immunostaining results and prior measurements support the presence of both H3K27me2 and H3K27me3 on maternal chromosomes (Fig. 4A, D) (18,23). We chose the aforementioned proportions of methylation groups as an initial condition, as they serve as a conservative estimate of the maternal methylation status. We did not initialize maternal chromosomes with any H3K27me1, as we were unable to measure this mark with immunostaining due to the incompatibility of the available antibodies with the immunostaining protocol. The positions of the methylation groups along the initialized chromatin fiber were randomly determined. To stimulate paternally-provided chromatin, a chromatin fiber was initialized with 100% of its H3K27 positions with me0, as it has been reported that paternal chromosomes lack all forms of H3K27 methylation (23).

#### 1.3 Nuclear cycle timing

Cell cycle timing in early *Drosophila* development is reproducible and well characterized. Our algorithm incorporates the following cell cycle durations approximated from our measurements and prior reports (Table S1) (54–56).

#### 1.4 DNA replication

We specified in the model that DNA replication occurs four minutes following each mitosis, consistent with prior estimates that DNA replication begins around that time (55,57). As modeled previously, the algorithm accounts for DNA replication in a single time step, rather than progressively over a cell cycle (25). To simulate DNA replication, the algorithm reduces both H3K27 positions at each nucleosome to me0 with a probability of 0.5.

The model does not incorporate sources of demethylation beyond DNA replication, given prior modeling work shows a baseline level of demethylation negatively impacts how well their model fits experimental data.

### 2. Nuclear E(z) and Pho concentrations

As our prior live-imaging measurements were restricted to NC11-NC14 (18), we developed models to describe the nuclear dynamics of E(z) and Pho throughout the first 13 cleavage cycles and the three subsequent cell cycles of development. To estimate E(z) levels on a per-cell cycle basis, we fit an equation for exponential decay to the measured nuclear EGFP-E(z) fluorescence at the center time points of NC11, NC12, NC13, and NC14 (Fig. S1 A). The fit suggests that, with each cell division, nuclear E(z) concentrations decrease by 24% (Fig. S1 A). Given our prior immunostaining measurements however, we expect nuclear E(z) concentrations to be relatively constant before NC11 (18). Therefore, we modeled nuclear E(z) concentrations as constant from NC1 to NC10, holding the concentration at the value predicted for NC10 (Fig. S1 B). Following NC10, the model allows nuclear EGFP-E(z) to decay exponentially on a per-cell cycle basis according to the exponential decay equation (Fig. S1 A-B). In the model, nuclear E(z) concentrations at all time points in a cell cycle following NC9 were determined based on the prediction the exponential decay equation makes for the cell cycle (Fig. S1 A-B). To predict nuclear Pho levels, we fit a sigmoidal equation to the live-imaging measurements at the center time points of NC11, NC12, NC13, and NC14 (Fig. S1 C). We find our measurements follow sigmoidal dynamics well (Fig. S1 C). To model nuclear Pho after NC14, we held Pho constant at its predicted NC14 value from NC14 onward; we figured we cannot assume that Pho continues to increase in nuclei after the last time point of our measurements. To model nuclear Pho concentrations at and prior to NC14, we used the per-cell cycle values produced by the sigmoidal equation (Fig. S1 C-D). At all time points across a cell cycle prior to the 15th cycle, the model holds nuclear Pho levels at the value the sigmoidal equation predicts for the cell cycle (Fig. S1 C-D). For convenience, we did not account for the mitotic periods when E(z) and Pho are absent from nuclei.

To better incorporate Pho and E(z) concentrations into the stochastic model of H3K27 methylation, we sought to convert the concentration estimates of normalized fluorescence units (AU) into nM. Therefore, we drew upon published mass spectrometry measurements of E(z) and Pho levels in 2- 4 hour *Drosophila* embryos (44). We converted the published values of an average of 3,300 E(z) molecules and 9,808 Pho molecules per 1C genome into nM nuclear concentration estimates using the average volume of a nucleus during that time in development, which we calculated assuming a nucleus is a sphere with a diameter of 7 μm (the average diameter we determined from our live-imaging data) (44). Our calculations produced a nuclear E(z) concentration estimate of 61 nM and a nuclear Pho concentration estimate of 181 nM for 2-4 hr, or approximately NC14, embryos. To extrapolate nM concentrations in the proceeding and subsequent cell cycles from these values, we normalized the modeled E(z) and Pho dynamics by their predictions for NC14, and then multiplied the normalized predictions by 61 nM and 181 nM for the E(z) and Pho models respectively. We incorporated the modeled E(z) and Pho nM concentration dynamics into our “Dynamic Pho” stochastic simulations of H3K27 methylation (Fig. 2B, D-E). For our “Constant Pho” simulations, Pho was held constant at its maximal concentration of 181 nM while E(z) concentrations were varied according to the described dynamics.

### 3. E(z) reaction probabilities

The efficiency with which E(z) catalyzes a methylation reaction varies depending on its substrate (39). It has been reported that the efficiencies for the me0-me1, me1-me2, and me2-me3 reactions follow a 13:4:1 ratio, with the me0-me1 reaction occurring with the highest efficiency and the me2-me3 occurring with the lowest (39). Like prior work, for each H3K27 position, the model therefore scales the reaction probability of E(z) according to the methylation state of the position in question: the ratio *P*_0-1_: *P*_1-2_:*P*_2-3_ is 13:4:1 (25). Our model incorporates a baseline methylation probability for the me2-me3 reaction like the prior model, which we refer to as *V* (25). Whether a potential reaction is driven by allostery or nucleation, along with the starting state of the H3K27 position, determines how *V* is scaled at that position (Section 3.2: Allostery, Section 3.3: Nucleation).

#### 3.1 E(z) and Pho binding

In the model, the probability that E(z) or both E(z) and Pho occupy the nucleosome array contributes to the methylation reaction probability. The model calculates the probability of E(z) binding to a site within the domain from the E(z) concentration, *[E(z)]*, and the dissociation constant of the E(z)-chromatin interaction, *K_e_*. Given the statistical weight of the E(z) occupied state, *e* = *[E(z)]*/*K_e_*, the probability of E(z) occupying the domain, *y_e_*, is

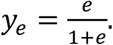

Likewise, the model calculates the probability of Pho binding to a site within the domain, *y_p_*, using the statistical weight of the Pho occupied state, *p* = *[Pho]*/*K_p_*, which depends on the nuclear Pho concentration, *[Pho]*, and the the dissociation constant of the Pho-chromatin interaction, *K_p_*. Therefore,

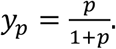

Similarly, the model calculates the probability that both E(z) and Pho occupy the domain, *y_ep_*, with

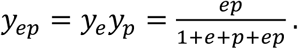

These probabilities are incorporated into the mechanisms of allostery and nucleation as described below.

#### 3.2 Allostery

The presence of H3K27me/3 at neighboring nucleosomes increases the catalytic rate of E(z) through an allosteric mechanism (30). For a given H3K27 position, our algorithm checks the methylation status of all four of the H3K27 positions on neighboring nucleosomes. If at least one of the neighbors is me2/3, the model scales the methylation probability of E(z) by the allosteric constant used in prior work, *allo* = 3.5 (25). This implementation of allosteric stimulation differs from prior work, as the model of Lundkvist et al. randomly chooses a partner histone to check if there is me3 and therefore allosteric stimulation (25). By allowing for any of the neighboring histones to stimulate a reaction, and for me2 in addition to me3 to stimulate E(z), our model amplifies the effects of allostery on the methylation dynamics at a locus. The model uses the following expressions to determine the probability of adding a methylation group, *k*, to a given H3K27 position (Table S2).

To decide whether E(z) will methylate, the algorithm draws from Bin(1, *k*).

#### 3.3 Nucleation

If a methylation reaction does not occur at a H3K27 position due to allosteric stimulation, the model provides E(z) another opportunity to catalyze a reaction if both E(z) and Pho simultaneously occupy the locus. In a nucleation-driven reaction, *allo* = 1, therefore negating the contribution of allostery, and the methylation probabilities depend on *y_ep_*. The model uses the following expressions to determine the probability of adding a methylation group, *k*, to a given H3K27 position due to nucleation (Table S3).

As in the case of an allosteric-driven reaction, the algorithm draws from Bin(1, *k*) to decide whether E(z) will methylate.

#### 3.4 E(z) clustering

With the baseline methylation probability we chose for our simulations, the algorithm predicts that multiple methylation reactions occur in each time step (Section 4: Parameter calibration, Table S4). Between maternal and paternal simulations, on average 4.2 reactions occur every 15 seconds. Given reported catalytic rates of E(z), it is extremely unlikely that a single E(z) molecule could catalyze multiple reactions in such a short interval (36). Therefore, we propose that clusters of E(z) molecules act at PcG domains, allowing for methylation of different histones simultaneously. *In vitro* measurements indicate that, at most, the turnover of PRC2 after catalyzing me0-me1 is approximately 1 min^-1^ (38–40). Therefore, we expect that at a minimum, a cluster contains 4.2 x 4 = 17 E(z) molecules on average.

### 4. Parameter calibration

Our model incorporates the following parameters: an allosteric stimulation constant (*allo*), a baseline me2-me3 reaction probability (*V*), a dissociation constant for the E(z)-chromatin interaction (*K_e_*), and a dissociation constant for the Pho-chromatin interaction (*K_p_*). Our model uses *allo* = 3.5, the best-fit parameter determined in prior work (25). For the dissociation constants of E(z) (*K_e_*) and Pho (*K_p_*), the model uses equivalent values, as chromatin binding rate constants for these factors determined from prior FCS experiments have been reported to be of the same order of magnitude (58). We specified *K_e_* = *K_p_* = 10 nM to match approximately the affinity of the PRC2 complex to a nucleosome array measured *in vitro* (59). Given these decisions, the only free parameter left in this model was *V*, the baseline me2-me3 methylation probability. We determined the value of *V* that allowed the model to best predict our H3K27me3 ChIP-seq measurements over NC14 by performing a parameter sweep, the results of which are presented in the main text (Fig. 2C). For the parameter sweep, we used the maternal initial conditions of the model, where each chromatin fiber was initialized with 50% of its H3K27 positions with me0, 0% of its positions with me1, 25% of its positions with me2, and 25% of its positions with me3.

#### 4.1 H3K27me3 at isolated E(z) peaks

To evaluate the model’s performance, we compared simulation outputs to H3K27me3 ChIP-seq signal across 54-kilobase regions each containing an isolated E(z) peak (Fig. 1B). E(z) peaks were chosen for analysis if they occupy an H3K27me3 domain, if they are in the top 90% of E(z) peaks in terms of max signal, and if they are more than 27 kilobases (or 150 nucleosomes, at 180 bp/nucleosome) away from neighboring E(z) peaks. We selected the H3K27me3 profiles at NC14- early, NC14-mid, and NC14-late timepoints over 27-kilobase segments on either side of the chosen E(z) peaks. By selecting H3K27me3 regions this way, we isolated the activities of single PREs on H3K27me3 dynamics. We compared simulation results to the mean profiles of H3K27me3 at these peaks (Fig. 1B). To allow for comparison with our simulations, we integrated the H3K27me3 profiles at each time point and normalized the integrated values, finding normalized peak integrations of 0.25, 0.52, and 1 for NC14-early, NC14-mid, and NC14-late time points respectively. As discussed in the main text, with a parameter sweep we determined that a baseline methylation probability of *V* = 1.075 x10^-3^ allows the model to most accurately simulate the relative increases in H3K27me3 at isolated PREs over NC14 (Fig. 2C).

### 5. Parameter summary

### 6. Drosophila stocks

The wild type stock used in this study is *w^1118^* as it serves as the genetic background for gene edited and transgenic lines in our laboratory. *E(z)^61^* mutant embryos were produced by culturing homozygous male and female *E(z)^61^* adults at 29°C overnight prior to embryo collection. *E(z)^61^/TM3* was a kind gift of Richard Jones, and is a temperature-sensitive loss-of-function mutant allele that is expressed at the restrictive temperature of 29°C (60). *sesame* mutant embryos were produced by collecting eggs laid by homozygous *sesame* mutant mothers crossed to male siblings. The *sesame^185b^* mutant stock was originally provided by Benjamin Loppin to the Princeton Stock Collection. Mutant cuticle phenotypes and lethality rates were measured prior to further experimentation to confirm that stocks bred true.

### 7. Immunostaining

#### 7.1 Fixation

Embryos were collected from flies of a genotype of interest (*w^1118^*, *E(z)^61^*, or *sesame^185b^*) after a laying period of 2-6 hours on an apple-juice agar plate. A heat-methanol fixation process was then performed as reported previously (18). Embryos were dechorionated for 1 minute in 4% Sodium Hypochlorite and then washed with deionized water. Following dechorionation, embryos were immersed in 5 ml of boiling 0.3% Triton X-100 + 0.4% (w/v) NaCl in a 20 ml scintillation vial for 10-20 seconds. The vial with the embryos was then placed on ice to cool for ≥ 15 minutes. Once ice-cold, the salt solution was removed (leaving the embryos). 5 ml methanol and 5 ml heptane were then added and the vial vortexed for 30 seconds to help the embryos dissociate (“pop”) from their vitelline membranes. The popped embryos that fell to the bottom of the vial were collected and washed 3x in methanol. The embryos were then stored at -20°C for ≥ 24 hours.

#### 7.2 Immunostaining

Fixed embryos were rehydrated in PBT (1x phosphate buffered saline + 0.1% Triton X-100 + 1% Bovine Serum Albumin) and then blocked for 1 hour in blocking buffer (1x phosphate buffered saline + 0.1% Triton X-100 + 5% Bovine Serum Albumin) rocking at room temperature. Embryos were then incubated in blocking buffer and primary antibody rocking overnight at 4°C. To stain H3K27me3, the anti-H3K27me3 antibody (Diagenode C15410195) was diluted 1:500 in blocking buffer. To stain H3K27me2, the anti-H3K27me2 antibody (Cell Signaling #9728) was diluted 1:1000 in blocking buffer. The specificities of the anti-H3K27me3 and me2 antibodies were previously tested using the methodology used here (61) and reported here: https://chromatinantibodies.com/. Staining of H3K27me1 was not feasible, as the available antibodies are incompatible with the immunostaining protocol. After overnight incubation, the primary antibody solution was removed and embryos washed in PBT. Following the washes, embryos were incubated in blocking buffer and secondary antibody (goat anti-rabbit coupled to Alexa 546 nm, 1:500 dilution) rocking for 2 hours at room temperature. The secondary antibody solution was then discarded and embryos rinsed in PTw (1x phosphate buffered saline + 0.1% Tween-20). Embryos were then stained with 1 μg/mL DAPI (4’,6-diamidino-2-phenylindole; Invitrogen REF: D1306) in PTw rocking for 1 hour at room temperature. The DAPI solution was replaced with PTw and the embryos stored at 4°C.

#### 7.3 Imaging

Slides were prepared for imaging by mounting embryos in Prolong Gold Antifade Mountant (Invitrogen P36934) and overlaying them with a glass coverslip. Images were collected as reported previously (18) on a Leica SP8 WLL Confocal Microscope using a 63x/1.3 NA glycerol- immersion objective. To create the presented time course images, staged embryos were imaged with a 100 Hz scan rate at 512x512 pixels,12-bit resolution, and 2x zoom. Z-stacks were imaged with a 1 μm z-step size and positioned such that the first and last slices of the stacks corresponded to the top and bottom of the nuclei. To create the *wild type* NC8 and *sesame* NC9 high-resolution images, embryos were imaged with a 100 Hz scan rate at 1024x1024 pixels,12-bit resolution, and 4x zoom. In this case, z-stacks were imaged with a 0.33 μm z-step size and positioned such that the first and last slices of the stacks corresponded to the top and bottom of the nuclei. A 557 nm laser was used to excite Alexa 546 nm conjugated secondary antibodies. A 405 nm laser was used to excite DAPI. Images were collected with a by-line sequential scan of the lasers.

## Author contributions

**E.A.D.:** original draft preparation (lead); review and editing (equal); conceptualization (equal); methodology (lead); investigation (lead); software (lead); formal analysis (lead); data curation (lead); visualization (equal). **N.G-S.:** conceptualization (equal); review and editing (supporting); visualization (supporting). **S.A.B.:** original draft preparation (supporting); supervision and administration (lead); funding acquisition (lead); conceptualization (equal); methodology (supporting); review and editing (equal); visualization (equal).

## Acknowledgments

We thank Eric Wieschaus for comments and critiques on an earlier version of this manuscript. We thank Richard Jones for generously providing the *E(z)^61^/TM3* stock. We thank the Bloomington Drosophila Stock Center for providing additional stocks and Flybase for providing an essential resource to the *Drosophila* community. E.A.D. was supported by the Cellular and Molecular Basis of Disease training program (T32 GM008061), is supported by an NSF GRFP fellowship (DGE-2234667), and is a Data Science Fellow at the Northwestern Institute on Complex Systems. Experiments were supported by the National Institutes of Health grant R01 HD101563 to S.A.B.. S.A.B. is a Pew Scholar in the Biomedical Sciences, supported by the Pew Charitable Trusts.

---

**Figure S1:**
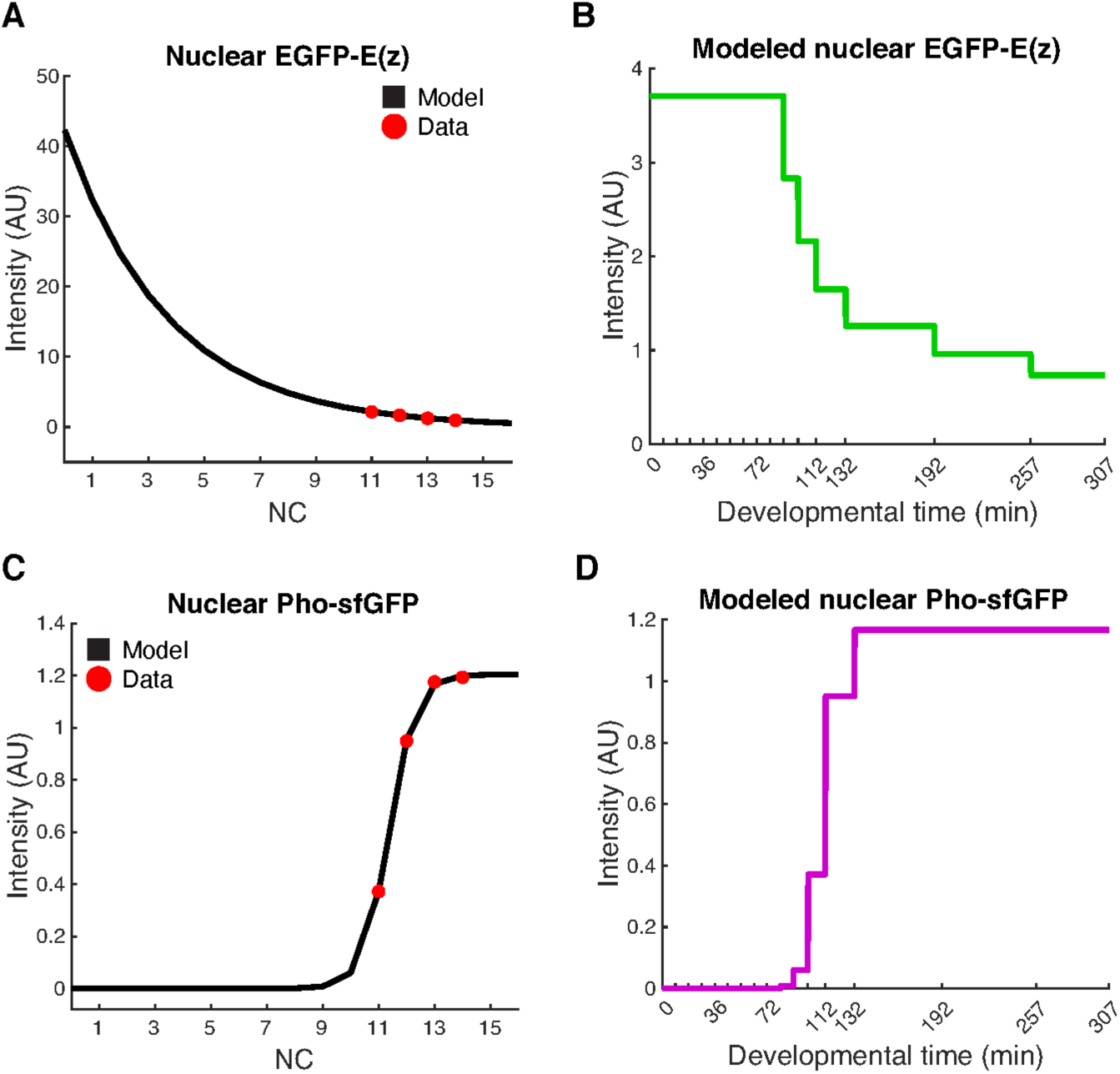
Models of E(z) and Pho nuclear dynamics. **A)** Shown are the normalized nuclear fluorescence measurements of EGFP-E(z) at the center points of NC11, NC12, NC13, and NC14 (red dots). The exponential decay equation *y* = 42.3(0.7631)*^x^* predicts the nuclear EGFP-E(z) intensity, *y*, for the *x* cell cycle (black line). **B)** Shown is a model of nuclear EGFP-E(z) that describes fluorescence intensities across the first 16 cell cycles of development. E(z) levels are held constant NC1-NC10, and then decrease with each subsequent cell cycle according to the values specified by the exponential decay equation in (A). **C)** Shown are the normalized nuclear fluorescence measurements of Pho-sfGFP at the center points of NC11, NC12, NC13, and NC14 (red dots). The sigmoidal equation *y* = 1.204/(1+exp(-2.135*(*x*-11.38))) predicts the nuclear Pho-sfGFP intensity, *y*, for the *x* cell cycle (black line). **D)** Shown is a model of nuclear Pho-sfGFP that describes fluorescence intensities across the first 16 cell cycles of development. Pho levels increase with each cell cycle according to the values specified by the sigmoidal equation in (C), and then are held constant following NC14.

**Table S1:**
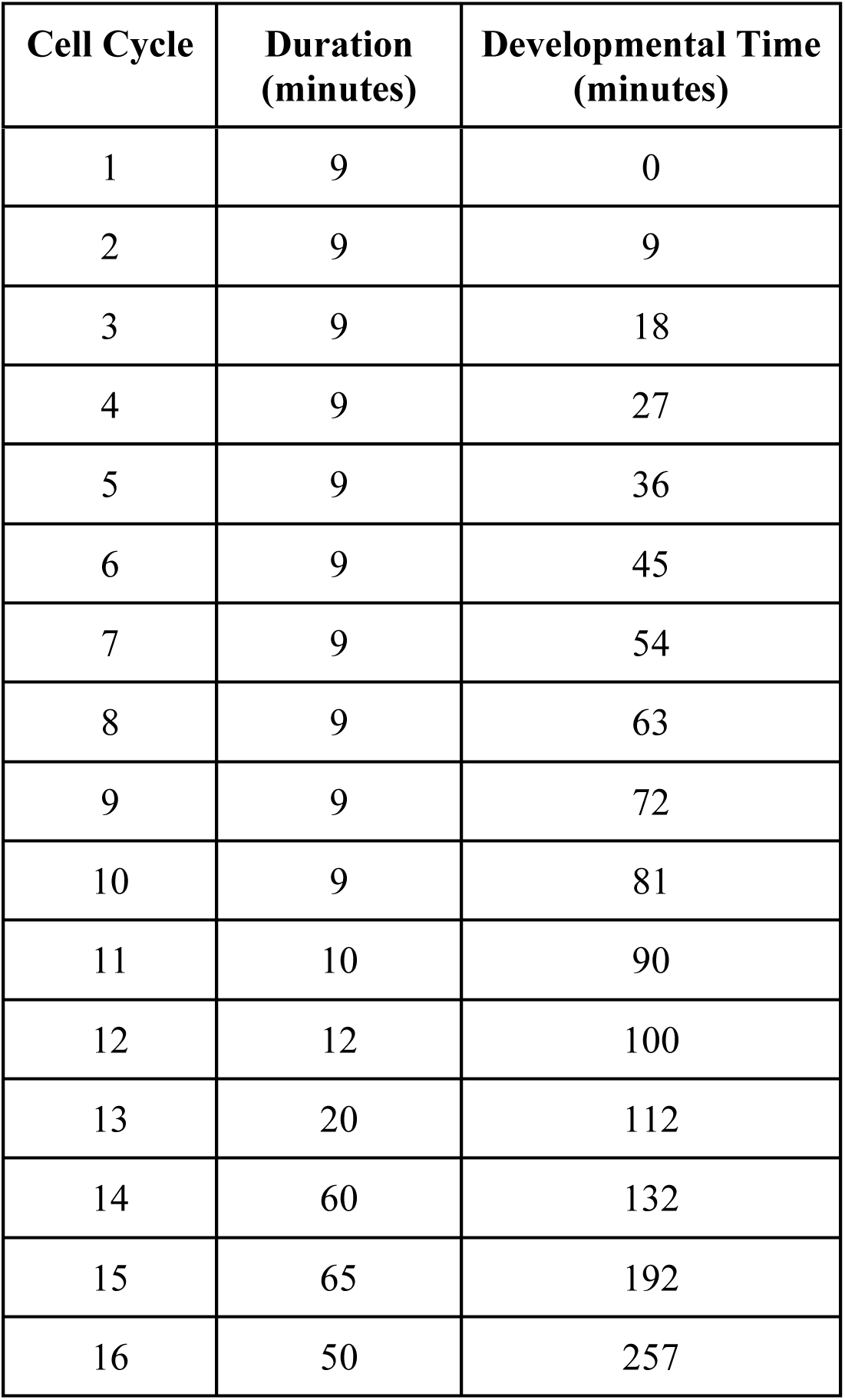
Cell cycle timing used in simulations. Shown are the durations (center column) of the first 16 cell cycles of *Drosophila* development (left column). The right column reports the developmental time points corresponding to the start of each cell cycle. Values are estimated from a combination of prior reports and live-imaging measurements (54–56).

**Table S2:** H3K27 methylation reaction probabilities for the allosteric mechanism. The model scales methylation probabilities based on the starting methylation state of the H3K27 position in question (left column) and the 13:4:1 ratios of H3K27me reaction efficiencies reported in prior work (right column) (39). In the case of allosterically-driven E(z) activity, the model also scales methylation probabilities by the allosteric stimulation parameter, *allo* = 3.5, and the probability that E(z) occupies the modeled nucleosome array, *y_e_* (right column).

| H3K27 Methylation State | Reaction Probability, $k$ |
| --- | --- |
| me<sub>0</sub> | $13 * V * allo * y_e$ |
| me<sub>1</sub> | $4 * V * allo * y_e$ |
| me<sub>2</sub> | $V * allo * y_e$ |
| me<sub>3</sub> | 0 |

**Table S3:**
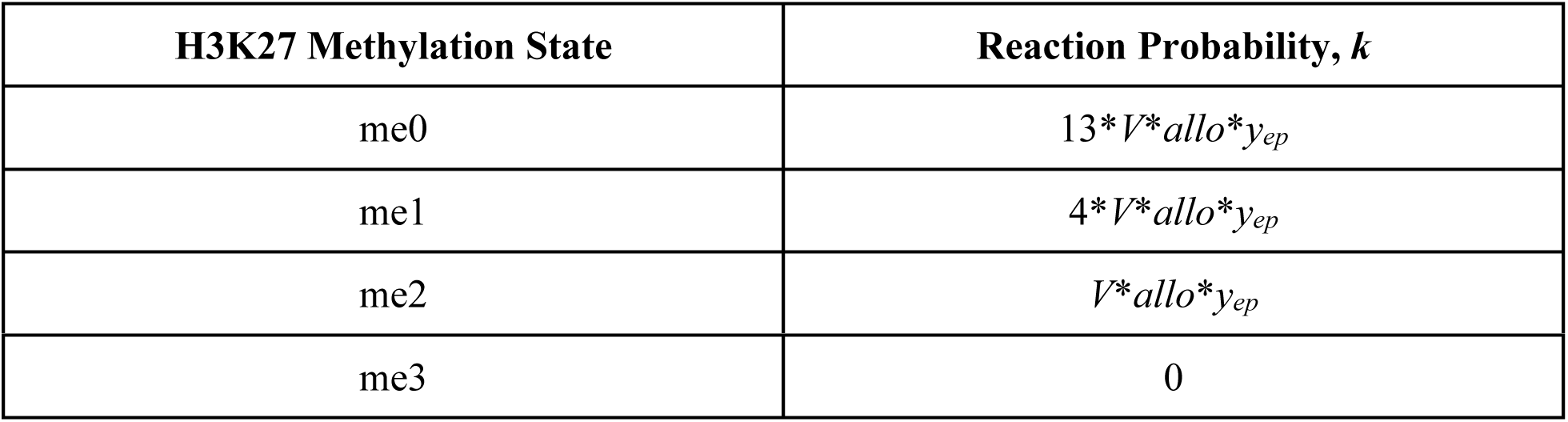
H3K27 methylation reaction probabilities for the nucleation mechanism. As in the allosteric mechanism, the model scales methylation probabilities based on the starting methylation state of the H3K27 position in question (left column) and the 13:4:1 ratios of H3K27me reaction efficiencies reported in prior work (right column) (39). For nucleation-driven E(z) activity, the model also scales methylation probabilities by the probability that E(z) and Pho simultaneously occupy the modeled nucleosome array, *y_ep_* (right column). While still included in the equations (right column), the allosteric stimulation parameter does not impact methylation probabilities in the case of nucleation, as *allo* = 1.

**Table S4:**
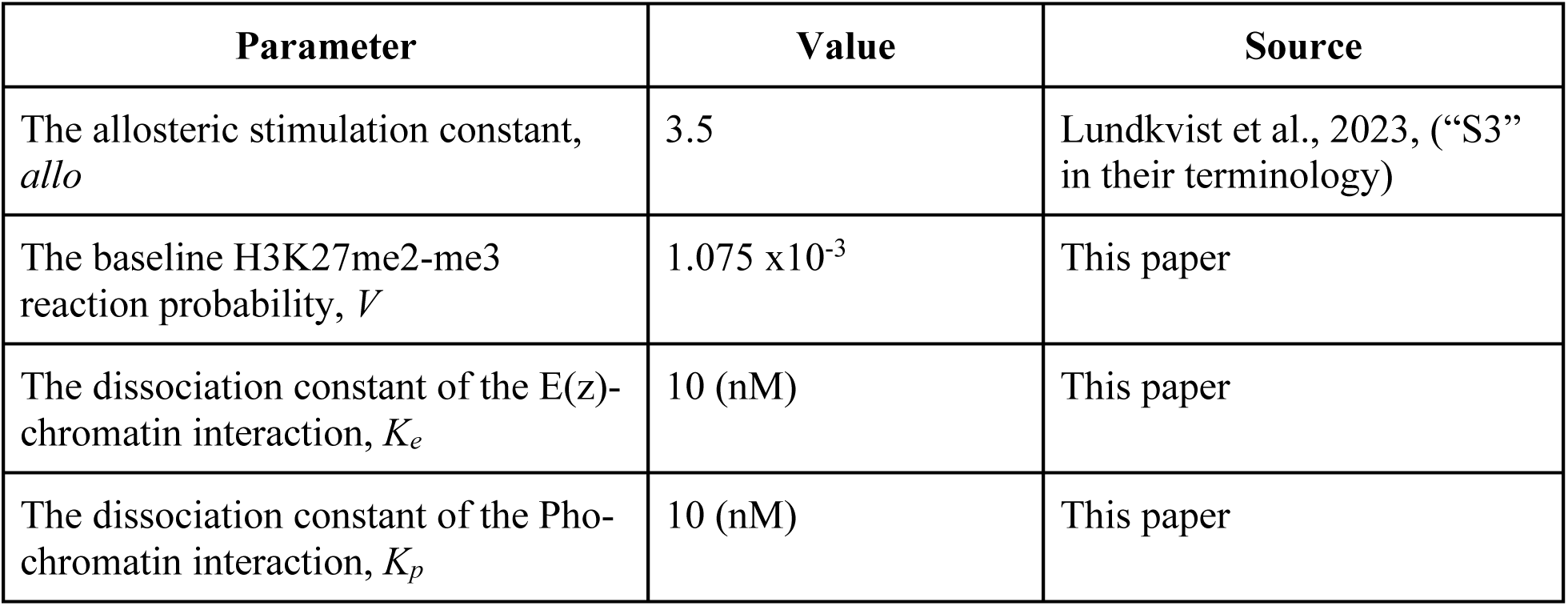
Parameter values used for modeling H3K27 methylation.

| Parameter | Value | Source |
| --- | --- | --- |
| The allosteric stimulation constant, *allo* | 3.5 | Lundkvist et al., 2023, (“S3” in their terminology) |
| The baseline H3K27me2-me3 reaction probability, *V* | $1.075 \times 10^{-3}$ | This paper |
| The dissociation constant of the E(z)-chromatin interaction, *K<sub>e</sub>* | 10 (nM) | This paper |
| The dissociation constant of the Pho-chromatin interaction, *K<sub>p</sub>* | 10 (nM) | This paper |

